# Ion occupancy of the selectivity filter controls opening of a cytoplasmic gate in the K2P channel TALK-2

**DOI:** 10.1101/2023.11.22.568211

**Authors:** Lea C. Neelsen, Elena B. Riel, Susanne Rinné, Freya-Rebecca Schmid, Björn C. Jürs, Mauricio Bedoya, Jan P. Langer, Bisher Eymsh, Aytug K. Kiper, Sönke Cordeiro, Niels Decher, Thomas Baukrowitz, Marcus Schewe

## Abstract

Two-pore domain K^+^ (K_2P_) channel activity was previously thought to be controlled primarily via a selectivity filter (SF) gate. However, recent crystal structures of TASK-1 and TASK-2 revealed a lower gate at the cytoplasmic pore entrance. Here, we report functional evidence of such a lower gate in the K_2P_ channel K2P17.1 (TALK-2, TASK-4). We identified compounds (drugs and lipids) and mutations that opened the lower gate allowing the fast modification of pore cysteine residues. Surprisingly, stimuli that exclusively target the SF gate (i.e., pH_e_., Rb^+^ permeation, membrane depolarization) also opened the cytoplasmic gate suggesting that the SF can induce global structural changes in TALK-2. Reciprocally, opening of the lower gate reduced the electrical work required to force ions into the SF to induce its opening as apparent in large shifts of the conductance-voltage (G-V) curves. These shifts, thereby, represent the mechanical work done by the SF to induce a global structural re-arrangement that opened the lower gate. In conclusion, it appears that the SF is so rigidly locked into the TALK-2 protein structure that changes in ion occupancy can pry open a distant lower gate. Vice versa, we show that opening of the lower gate concurrently forces the SF gate to open. This concept might extent to other K^+^ channels that contain two gates (e.g., voltage-gated K^+^ channels) for which such a positive gate coupling has been suggested, but so far not directly demonstrated.

**Synopsis:**

- TALK-2 channels, like most K_2P_ channels, possess a functional gate in the selectivity filter (SF; the upper gate) that is opened by rising extracellular pH and voltage-dependent ion binding (voltage gating).
- A second (lower) permeation gate in TALK-2 at the cytoplasmic end of TM4 is identified using cysteine modification, scanning mutagenesis and structural modelling. This gate can be opened by anionic lipids (LC-CoA) as well as pharmacological ligands (e.g., 2-APB).
- The modification reactivity of a cysteine introduced between the two gates reveal that stimuli targeting the SF gate also open the lower gate. Furthermore, stimuli that open the lower gate reduce the voltage (i.e., electrical work or mechanical load) required to open the SF gate. These findings demonstrate a tight positive coupling between the two gates.
- The concept of strong positive gate coupling might extend to other K^+^ channels with two gates (e.g., voltage-gated K^+^ channels) for which positive gate coupling has been suggested but so far not directly demonstrated.

## Introduction

TWIK-related alkaline-pH-activated potassium (TALK-2, K_2P_17.1, *KCNK17*) channels are members of the two-pore domain K^+^ (K_2P_) channel family. They were also referred as TWIK-related acid-sensitive potassium (TASK-4) channels when first identified in 2001^1^. TALK-2 channels are expressed in various human cell types and organs (i.e., pancreas, aorta, brain, liver, placenta and heart) of the human body^1–3^. Despite their widespread distribution in several tissues the functional role of these channels in biological processes has not been established yet, especially in contrast to other well-investigated acid-sensitive K_2P_ channels such as TASK-1^4–9^. However, its malfunction, up- or down-regulation or genetic variants of TALK-2 K_2P_ channels have been associated with a number of cardiovascular diseases such as cardiac conduction disorders^10^, ischemic stroke and cerebral haemorrhage^11–14^ as well as arrhythmias including atrial fibrillation, idiopathic ventricular fibrillation and long QT syndrome^10,15,16^. Furthermore, thus TALK-2 channels are highly and specifically expressed in the human pancreas and are considered as a risk factor for the pathogenesis of type 2 diabetes^17,18^.

TALK-2 channels are activated by alkaline extracellular pH (pH_e_ > 7.4) that is thought to occur by deprotonation of an extracellular lysine causing the opening of the SF gate^1,2,18,19^. Furthermore, TALK-2 channel currents are enhanced by the production of nitric oxide radicals and reactive oxygen species^18^. Like most K_2P_ channels, TALK-2 channels are sensitive to changes in membrane voltage and permeating ion species^20^. Recently, polyanionic lipids of the fatty acid metabolism (e.g. oleoyl-CoA) have been identified as natural TALK-2 channel ligands, increasing channel activity by more than 100-fold^21^. How exactly these stimuli regulate the opening and closing of TALK-2 K_2P_ channels is, with the exception of voltage and pH_e_ acting at the SF gate, so far unclear.

TALK-2 channels are functional dimers and exhibit – as all other 14 K_2P_ channel family members – a characteristic topology of four transmembrane domains (TM1 to TM4) and two pore-forming domains (P1 and P2) within each channel subunit. The two P1 and P2 domains form the pseudo-tetrameric selectivity filter (SF) of the channel upon dimerization^22–24^. For the last two decades, K_2P_ channels were thought to be gated at the SF and, thus, that the various physicochemical stimuli acting on different regions of the channel (in particular on the cytoplasmic C-terminus) finally converge on the primary filter gate^20,25–29^.

Surprisingly, the recently resolved structures of TASK-1 and TASK-2, identified additional inner/cytoplasmic gates (further referred to as ‘lower’ gates)^30,31^. In TASK-1 this gate is formed by the crossing (therefore termed ‘X-gate’) of the late TM4 domains^31^, however, an activation mechanism for this gate is currently unknown. Based on the two cryo-EM structures of TASK-2 generated at pH 8.5 (open channel) and pH 6.5 (closed channel) the lower gate of TASK-2 is mainly formed by the interaction of two corresponding TM4 residues (K245, N243), hypothesised to function as molecular barrier in the process of pH gating^30^.

In this study we employed cysteine modification, alanine scanning mutagenesis homology modelling and various pore blocker to identify and characterize an additional lower gate in TALK-2 channels. These approaches provided information on the status of the SF gate and lower gate, respectively, and revealed that the two gates are strongly positively coupled. We show that the ion occupancy of the SF controls opening of the lower gate and estimated the electrical work required to open the lower gate. Our results establish a strong positive coupling of the two gates that can be envisioned as a concerted structural change involving both gates. Further, we demonstrate that the lower gate produced a state-dependent blocker pharmacology that is unique in K_2P_ channels.

## Results

### Probing for a cytoplasmic constriction in TALK-2 channels with cysteine modification

The basal activity of TALK-2 channels in excised patches is low, but pharmacological compounds (e.g. 2-APB)^32^ or polyanionic lipids (e.g. oleoyl-CoA)^21^ can induce large TALK-2 channel currents (**Fig. 1a, f**). However, the binding sites of these compounds and the mechanisms how they open the ion permeation pathway are currently unknown. To investigate the latter, we utilized a cysteine modification assay. Control experiments ensured that WT TALK-2 currents were insensitive to the application of the sulfhydryl reactive compound (2-(Trimethylammonium)ethyl) MethaneThioSulfonate (MTS-ET) regardless whether applied on the low activity basal state (**Supplementary Fig. 1a, Supplementary Table 1**) or the high activity state induced with 2-APB or oleoyl-CoA (**Supplementary Fig. 1c**). To test for a cytoplasmic constriction in TALK-2, we introduced a cysteine at amino acid position 145 (L145C) in TM2, that corresponds to a residue in TREK-1 (G186C) located in close proximity underneath the SF previously shown to result in a permeation block upon cysteine modification in TREK-1^29^. TALK-2 L145C mutant channels showed low basal activity, and both 2-APB and oleoyl-CoA produced robust activation very similar to the WT (**Fig. 1g, Supplementary Fig. 1d and Supplementary Fig. 5a**). Application of MTS-ET on the low activity basal state had no effect on the channel activity (**Fig. 1d left panel, Supplementary Fig. 1b**). In marked contrast, application of MTS-ET on L145C TALK-2 currents activated by 2-APB or oleoyl-CoA resulted in complete and irreversible current inhibition, indicating the chemical modification of L145C (**Fig. 1d middle and right panel, Supplementary Table 1**). These findings indicate that access of MTS-ET to L145C is blocked in the low-activity state of the channel but possible upon activation. To gain a better structural understanding, we generated TALK-2 homology models based on the TASK-1 and TASK-2 structures that both show a lower permeation constriction (see methods)^30,31^. In these models, the side chain of L145 points into the permeation pathway at a position between the SF and the lower gates (**Fig. 1b, c, Supplementary Fig. 2**). Furthermore, we used these TALK-2 models to pick a residue for cysteine substitution (Q266) pointing into the cytosol directly below the lower constrictions. Application of MTS-ET to Q266C TALK-2 mutant channels caused a mono-exponential and irreversible current activation (**Fig. 1e left panel**). Importantly, the observed modification occurred at a similar rate in both the low- and high-activity states suggesting similar access to the cysteine under both conditions (**Fig. 1e, Supplementary Table 2**). The fact that MTS-ET modification activated Q266C TALK-2 mutant channels might indicate that the modification at this position (i.e., close to the putative lower gate) destabilizes the closed gate and thus result in channel activation.

**Fig. 1.**
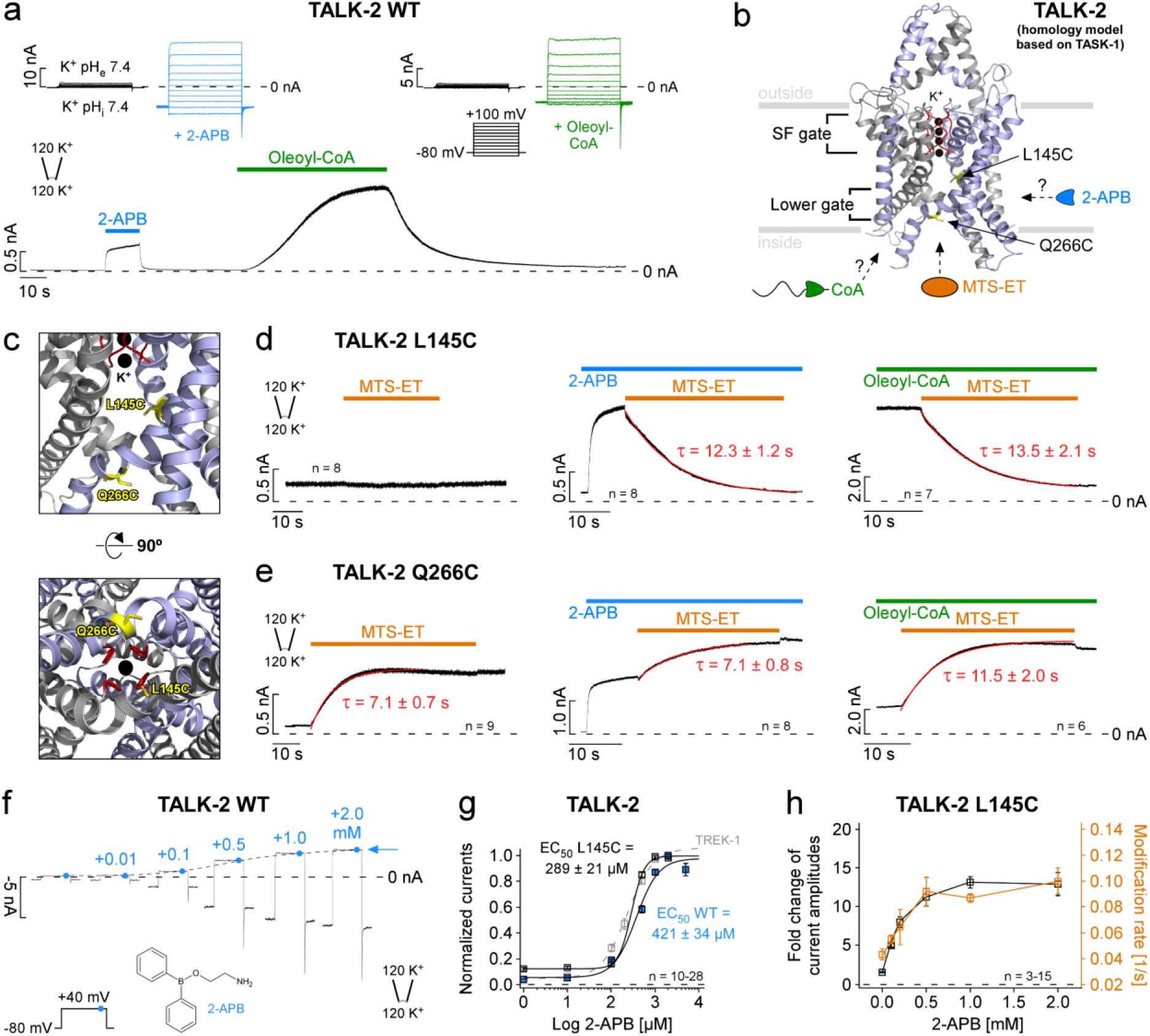
State-dependent modification of inner pore cysteine residues in TALK-2 K_2P_ channels. **a** Representative current trace measured under voltage-clamp conditions at +40 mV from inside-out patches of *Xenopus laevis* oocytes expressing WT TALK-2 channels symmetrical K^+^ (120 mM [K^+^]_ex._/120 mM [K^+^]_int._) at pH 7.4. Channel currents were activated with the indicated compounds (1 mM 2-APB and 5 µM oleoyl-CoA) applied to the intracellular side of the membrane. Inlays showing current-voltage responses of 2-APB (blue) and oleoyl-CoA (green) activation compared to the basal state (black) by the indicated voltage protocol. **b** Pore homology model of TALK-2 based on the crystal structure of TASK-1 (PDB ID: 6RV3, chains A, B) with the SF highlighted in red, K^+^ ions in black, and cysteine residues (L145C and Q266C) for MTS-ET modification in yellow. **c** Pore cavity zoom-in. Cut outs display the localization of L145C in the inner cavity and Q266C at the intracellular end of the pore. **d** Representative measurements of TALK-2 L145C channels showing state-dependent MTS-ET modification with no effect under unstimulated (basal) conditions or inhibition upon application of 1.0 mM MTS-ET in pre-activated states with 1.0 mM 2-APB (blue) or 5.0 µM oleoyl-CoA (green) with the indicated time constants (τ), respectively. **e** Same measurements as in (d) with TALK-2 Q266C channels showing state-independent modification with activation upon application of 1.0 mM MTS-ET. **f** Current responses recorded with the indicated voltage protocol in symmetrical K^+^ showing activation of WT TALK-2 with increasing 2-APB concentrations. The dotted line shows the increase and saturation of current amplitudes with 2-APB at + 40 mV. **g** 2-APB dose-response curves analyzed from measurements as in (f) for WT TALK-2 (blue), TALK-2 L145C (black) and WT TREK-1 (gray) channels. **h** Correlation between the fold change in current amplitudes of TALK-2 L145C channels at +40 mV (black squares) and the rate of MTS-ET modification (1/τ) at +40 mV (orange squares) with different 2-APB concentrations. Values are given as mean ± s.e.m with number (n) of experiments indicated in the figure and supplementary tables 1 and 2.

Our findings suggest the existence of a lower constriction blocking MTS-ET access at the level of the ‘X-gate’ in TASK-1 that is opened by 2-APB or oleoyl-CoA in TALK-2 channels. Accordingly, we observed that current activation and the rate of L145C modification concurrently increased with the 2-APB concentration levelling off at a concentration of 2.0 mM 2-APB that caused maximal current activation (**Fig. 1f-h, Supplementary Table 1**).

### The lower constriction functions as a permeation gate

Our data strongly suggest a gated lower pore constriction site. However, whether this constriction is an actual permeation gate is still disputable, as 2-APB and oleoyl-CoA could have also opened the SF gate to cause activation. Furthermore, a constriction that blocks MTS-ET access may not necessarily block ion permeation. Our attempts to use silver ions (Ag^+^, which is much smaller than MTS-ET and similar in size to K^+^) to probe for a permeation gate were not conclusive as Ag^+^ also inhibited WT TALK-2 channels. Therefore, we addressed this issue with a different approach by taking advantage of the fact that Rb^+^ has a strong activating effect on the TALK-2 SF^20^ and to do so, Rb^+^ needs to pass the lower constriction. And indeed, a stepwise increase in the 2-APB concentration resulted in a stepwise increase in Rb^+^ activation as if 2-APB activation had removed a constriction that prevented Rb^+^ to access the SF (**Fig. 2a, c**). Accordingly, the same effect on Rb^+^ activation was also observed when TALK-2 was activated by oleoyl-CoA (**Supplementary Fig. 3**). As a control, we performed the same experiment with TREK-1 channels, since these K_2P_ channels lack a lower gate but also undergo activation upon 2-APB application (**Fig. 1g**). In TREK-1, Rb^+^ activation was the strongest in the absence of 2-APB activation, indicating that Rb^+^ ions had free access to the SF in the low activity state of the channel (**Fig. 2b, c**). Moreover, 2-APB activation progressively covered up Rb^+^ activation consistent with the concept that TREK-1 channels lack a lower gate and both activating stimuli (2-APB and Rb^+^) converge onto the SF gate (**Fig. 2b**).

**Fig. 2.**
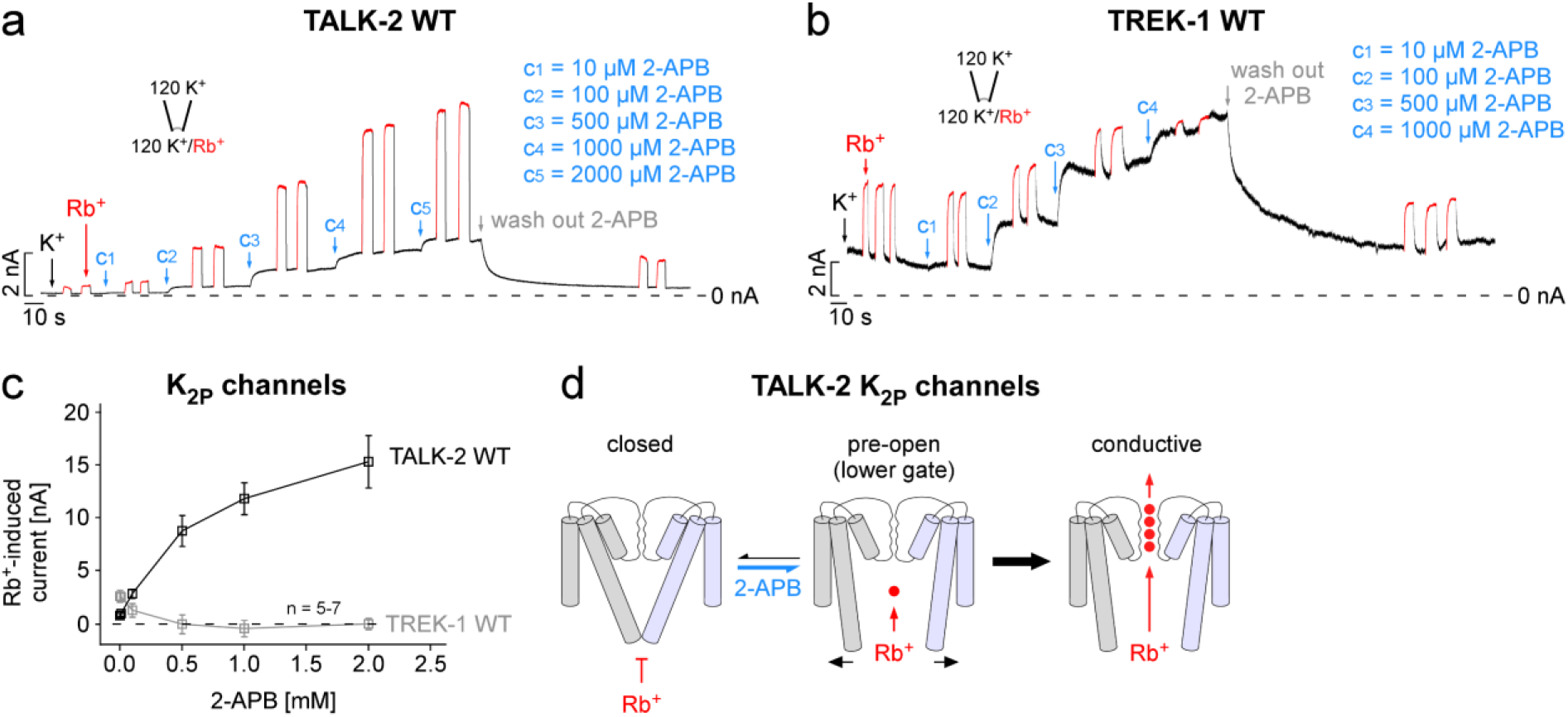
The lower constriction functions as a permeation gate. **a** Representative measurement of WT TALK-2 channels from an inside-out patch in symmetrical K^+^ at +40 mV with increasing 2-APB concentrations (c_1_ - c_5_) applied from the intracellular side at indicated time points (blue arrows). At steady-state current levels with 2-APB intracellular K^+^ was exchanged by Rb^+^ showing an enhanced activatory Rb^+^ ion effect on the SF in 2-APB pre-activated channels. **b** Same recording as in (a) for WT TREK-1 K_2P_ channels showing the stepwise loss of Rb^+^ activation in the presence of increasing 2-APB concentrations. **c** Correlation of Rb^+^-induced currents from measurements as in (a,b) in the presence of indicated 2-APB concentrations for either WT TALK-2 (black) or WT TREK-1 (gray) channels. **d** Gating scheme highlighting the effect of 2-APB and Rb^+^ on the lower and selectivity filter gate in TALK-2 channels. Values are given as mean ± s.e.m with number (n) of experiments indicated in the figure.

These results suggest that the lower constriction in TALK-2 channels functions as a permeation gate that must be open in addition to the SF gate to allow ion conduction (**Fig. 2d**).

### Characterization of the lower gate by scanning mutagenesis and single channel recordings

We performed systematic alanine scanning mutagenesis in the TM regions that contain the lower gates in TASK-1 and TASK-2 (i.e., TM2 and TM4) to identify mutations that would affect the stability of the lower gate in TALK-2 (**Fig. 3a, b, Supplementary Fig. 4a**). This approach identified four gain of function (g-o-f) mutations (V146A in TM2, W255A, L262A and L264A in TM4) that markedly increased TALK-2 basal currents 31.9 ± 3.4-fold (V146A), 4,5 ± 0.2-fold (W255A), 25.1 ± 2.1-fold (L262A) and 93.3 ± 9.8-fold (L264A) as assessed in two-electrode voltage-clamp (TEVC) experiments with *Xenopus* oocytes (**Fig. 3a**). Mapping of the g-o-f residues on our TALK-2 homology models showed a clustering of the g-o-f residues in the region of the lower pore constrictions identified in TASK-1 and TASK-2 at the cytosolic pore entrance of the channels (**Fig. 3c, Supplementary Fig. 4b**). In the TASK-1-based TALK-2 homology model the two residues showing the strongest g-o-f phenotype (i.e., V146A and L264A) are in direct proximity and could form a permeation constriction (**Fig. 3c**).

**Fig. 3.**
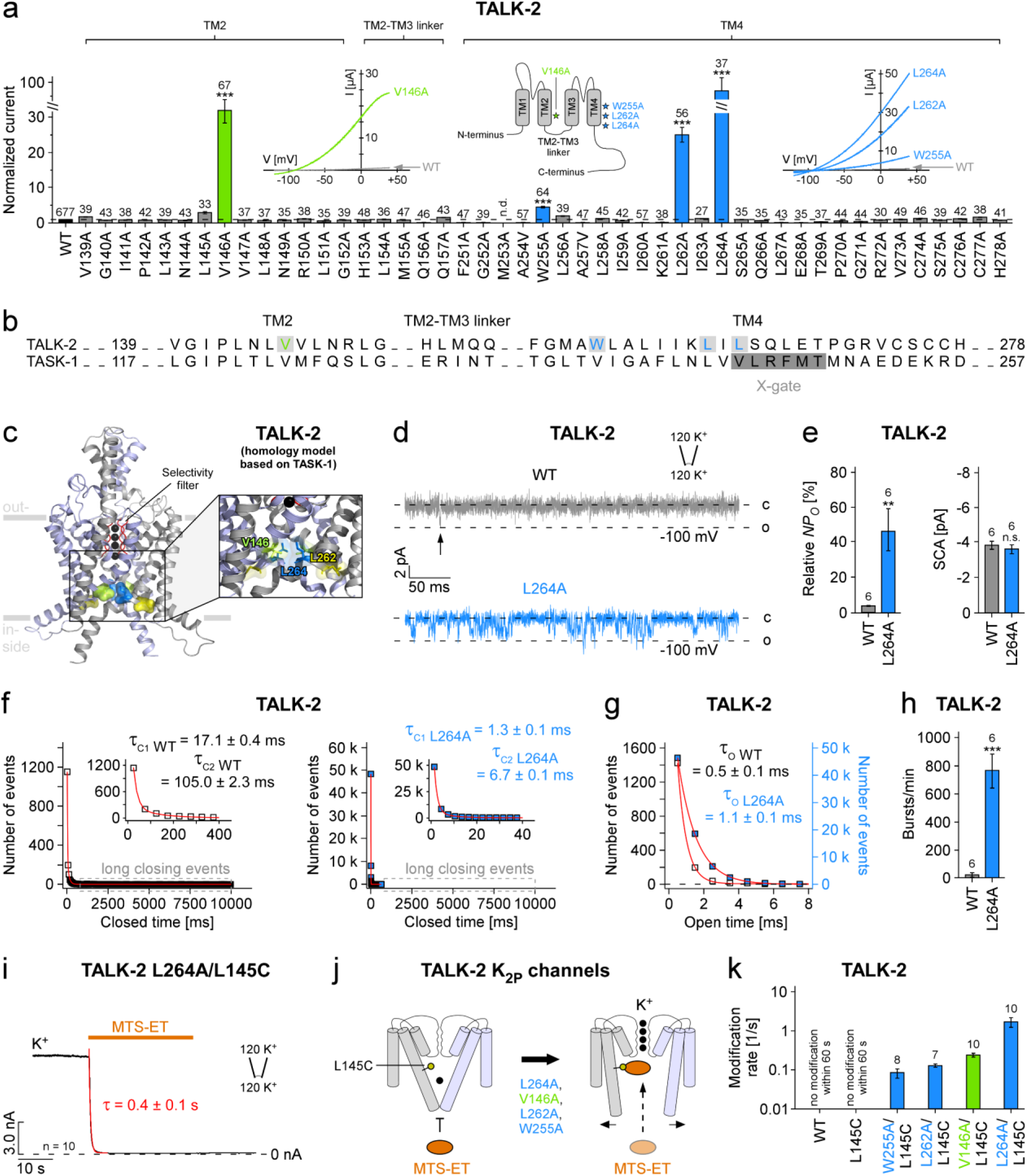
Functional characterization of the lower gate in TALK-2 K_2P_ channels. **a** Relative current amplitudes from two-electrode voltage-clamp (TEVC) measurements at pH 8.5 of WT and mutant TALK-2 channels 24 h after injection of 25 ng cRNA into *Xenopus laevis* oocytes. Currents were elucidated with a voltage protocol ramped from −120 mV to +45 mV within 3.5 s, analyzed at +40 mV and normalized to WT. Inlays showing representative WT TALK-2 (gray traces), TALK-2 L264A, L262A, W255A and V146A mutant channel currents (blue and green traces), respectively and a topology model of a channel protomer highlighting the localization of the g-o-f mutations in TM2 and TM4. **b** Sequence alignment of the TM2, TM2-TM3 linker, and TM4 regions of the human K_2P_ channels TALK-2 and TASK-1. **c** Pore homology model of TALK-2 based on the crystal structure of TASK-1 (PDB ID: 6RV3, chains A, B) highlighting the cluster of g-o-f mutations (V146A, L262A and L264A) at the cytosolic pore entrance. **d** Representative inside-out single channel measurements of WT TALK-2 (gray trace) and TALK-2 L264A mutant channels (blue trace) at −100 mV. **e** Relative open probability (*NP_O_*) and single channel amplitudes (SCA) analyzed from recordings as in (d) for WT and L264A TALK-2 channels (n = 6). **f-h** Analysis of the mean channel-open times (g), closed time events (f) and burst behavior (h; see methods) for WT and TALK-2 L264A mutant channels. TALK-2 L264A shows a complete loss of long closing events (area highlighted by dashed boxes) and decreased short (τ_C1_) and long (τ_C2_) closed times (f) and an increased bursting behavior (h) compared to WT (n = 6). **i** Representative measurement of TALK-2 L264A mutant channels additionally carrying the inner pore mutation L145C (TALK-2 L264A/L145C) at +40 mV showing a fast and irreversible modification and subsequent block upon application of 1.0 mM MTS-ET. **j** Cartoon illustrating the pore accessibility of MTS-ET in L145C mutant TALK-2 channels with or without carrying an additional g-o-f mutation. **k** Modification rates of WT, L145C and double mutant TALK-2 channels at +40 mV as indicated. Values are given as mean ± s.e.m with number (n) indicated above the bars in the figure and supplementary table 2 (n.d. = not determinable, n.s. = not significant, *** *P* ≤ 0.001, Student’s *t*-test).

WT and L264A mutant TALK-2 channels were further analyzed with inside-out single channel recordings (**Fig. 3d-h**). As expected, the basal single channel activity of WT TALK-2 was very low with a relative channel-open probability (*NP_O_*) of 3.9 ± 0.2 % and the L264A mutation resulted in a large increase to 46.7 ± 12.1 %, while the single channel amplitude was not affected (**Fig. 3d, e**). The increase in *NP_O_* appeared to primarily result from a destabilization of the closed state, as we observed a strong shortening of the short (∼13-fold) and long closed times (∼16-fold) (**Fig. 3f**), whereas, the open times were comparatively little (∼2-fold) affected (**Fig. 3g**). Overall, these changes result in a 29-fold increase of channel activity spend in a burst type mode with 767 ± 123 bursts/min for L264A channels compared to 27 ± 17 burst/min for WT channels (**Fig. 3h**). The observed shortening in closed times are consistent with the concept that the g-o-f mutations destabilize a permeation gate that now opens much more frequently. Further, if this destabilized permeation gate corresponds to the lower gate, we expect fast modification of the L145C mutation as indeed observed in L264A/L145C double mutant channels (**Fig. 3i, j**). Moreover, all four g-o-f mutants resulted in fast L145C modification (in the absence of ligand activation) (**Fig. 3i-k**) with rates that roughly correlated to the g-o-f effect (**Fig. 3a**) with L264A showing the fastest modification (**Fig. 3i, k, Supplementary Table 2**).

### Gate coupling in TALK-2 channels

The existence of two activation gates in TALK-2 raises the question if they are coupled. To address this question, we tested a stimulus that directly affects the SF gate. Extracellular alkalinization is thought to open the SF gate in TALK-2 by deprotonation of a lysine residue (K242) located at the outer pore helix connected to the SF^19^ (**Fig. 4b**). Accordingly, raising the pH_e_ from 7.4 to 9.5 resulted in large TALK-2 currents (**Fig. 4a, Supplementary Fig. 5a, b**). Under this condition we determined the MTS-ET accessibility of L145C and, surprisingly, observed a similar fast modification rate as seen with maximal (2.0 mM) 2-APB activation at pH_e_ 7.4 (**Fig. 4c, Supplementary Table 1**). However, raising the pH_e_ to 9.5 had little effect on the modification of Q266C consistent with the localization of this position at the cytoplasm-facing side of lower gate (**Supplementary Fig. 5c, Supplementary Table 2**). These results imply that opening the SF gate by high pH_e_ had also opened the lower gate in TALK-2.

**Fig. 4.**
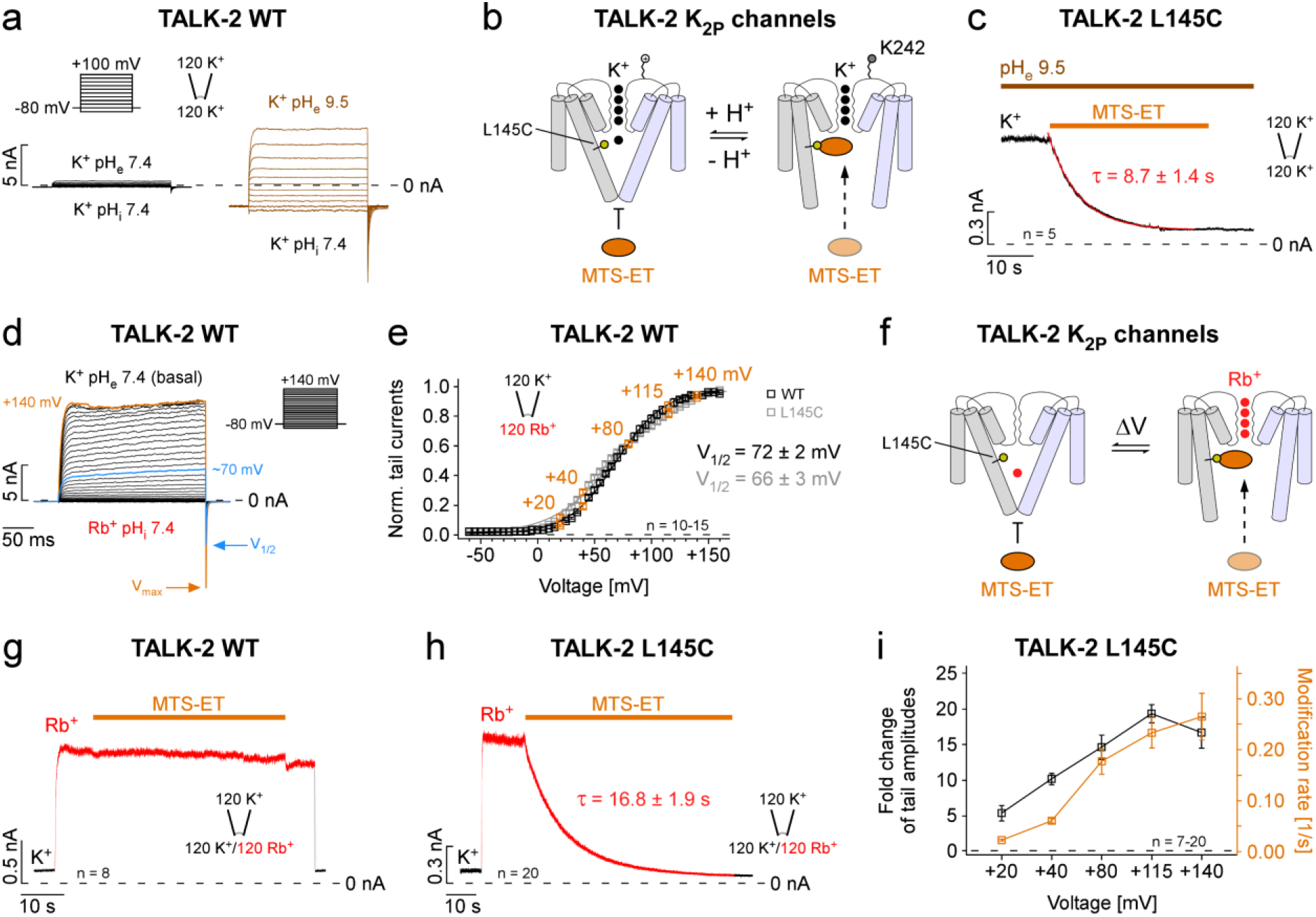
Direct stimulation of the SF produces a state-dependent cysteine modification in the pore of TALK-2. **a** TALK-2 current responses to voltage families as indicated in symmetrical K^+^ (120 mM [K^+^]_ex._/120 mM [K^+^]_int._) at pH 7.4 on both sides (black traces) and at extracellular pH 9.5 (brown traces). **b** Cartoon illustrating a simple TALK-2 channel gating model and pore accessibility to MTS-ET by alterations of the pH_e_ that directly affects the SF. **c** Representative modification and subsequent irreversible inhibition with 1.0 mM MTS-ET of TALK-2 L145C channels pre-activated by extracellular alkalinization (pH_e_ 9.5). **d** TALK-2 channel currents with intracellular Rb^+^ (120 mM [K^+^]_ex._/120 mM [Rb^+^]_int._) at pH 7.4 for different potentials as indicated showing a maximum *P_O_* reached for potentials positive to ∼+135 mV (V_max_), as further depolarizations do not increase the tail current amplitudes. **e** Voltage activation (conductance-voltage (G-V) curves) with V_1/2_ values of 72 ± 2 mV and 66 ± 3 mV of WT TALK-2 and L145C mutant channels, respectively. **f** Cartoon of a simple gating model with Rb^+^ as an amplifier for voltage activation targeting the SF and subsequently the lower gate in TALK-2 channels. **g**,**h** Representative measurements at +40 mV of WT (g) and L145C mutant TALK-2 channels (h) showing a non-modifiable state or almost complete modification/inhibition with 1.0 mM MTS-ET within 60 s in intracellular Rb^+^, respectively. **i** Correlation between the fold change of tail current amplitudes (black squares) of TALK-2 L145C channels and the incidental rate of MTS-ET modification (1/τ) (orange squares) with intracellular Rb^+^ at different potentials as indicated. Values are given as mean ± s.e.m with the number (n) of experiments indicated in the figure and supplementary table 1 - 3.

### Ion occupancy of the SF controls the opening of the lower gate

We have previously shown that the SF acts as an ion-flux voltage gate in many K_2P_ channels including TALK-2^20^. According to this concept, the low basal activity of exclusively SF-gated K_2P_ channels - such as TREK-1 - results from an inactivated and ion-depleted filter. Upon depolarization, ions are forced into the SF by the transmembrane electric field to induce voltage-dependent channel activation. The electrical work necessary to open the SF is reflected in the conductance-voltage (G-V) curves obtained by plotting the relative open probability (tail current amplitudes) against the membrane pre-pulse voltage (**Fig. 4d, e**). Particularly strong ion activation is seen with intracellular Rb^+^ as this ion appears to stabilize the conductive state of the filter more efficiently than K^+^ (**Fig. 4g, h, Supplementary Fig. 5d, e**). By monitoring the modification of L145C mutant channels we tested whether opening of the SF with voltage and Rb^+^ would also affect the status of the lower gate (**Fig. 4f,h and i**). Indeed, activation of TALK-2 channels upon K^+^ by Rb^+^ replacement at a membrane potential of +40 mV allowed strong L145C MTS-ET modification (**Fig. 4h, Supplementary Table 1**). Thus, the mere change in SF ion occupancy upon exchanging K^+^ by Rb^+^ in the SF is sufficient to markedly increase the *Po* of the lower gate allowing L145C modification (**Fig. 4f**). As expected, the L145C modification rate was strongly voltage-dependent and the increase in channel *P_O_* reflected in the G-V curve mirrored the increase in modification rate and, thus, voltage-dependent opening of the SF gate concurrently opened also the lower gate (**Fig. 4e, i**).

### The properties of the lower gate resulted in a state-dependent TALK-2 pharmacology

In voltage-gated K^+^ (K_v_) channels the closing of the lower gate can be prevented by the binding of blockers such as tetra-pentyl-ammonium (TPenA) in the pore cavity and, thereby, causing a slowing of the deactivation kinetics which is known as the ‘foot in the door’ effect^33,34^. To test for such a mechanism in TALK-2, we activated the channels with a voltage step to +100 mV in the presence of intracellular Rb^+^ followed by a repolarization step to −80 mV (to induce tail currents) with and without the K_2P_ channel pore blocker TPenA. Remarkably, we observed a tail current cross-over indicating that the deactivation time course is slowed by TPenA inhibition; i.e., with ∼50 % TPenA block of the tail current amplitudes, we observed a ∼2.5-fold slowing of the deactivation kinetics (**Fig. 5a**). We hypothesize, that the two gates are strongly coupled and, therefore, opening the SF gate by depolarization should also open the lower gate, which then consequently would allow blocker (like TPenA) binding within the pore cavity (**Fig. 5a cartoon**). Upon repolarization, however, the bound TPenA blocker obstructs closure of the lower gate and thereby, hinders the SF from closure/inactivation (strong gate coupling). In accordance, when we tested this protocol on TREK-2 K_2P_ channels, known to lack a lower gate, TPenA inhibition had no effect on the tail current kinetics indicating that here the SF gate is able to close unhindered with the pore blocker bound (**Fig. 5b**).

**Fig. 5.**
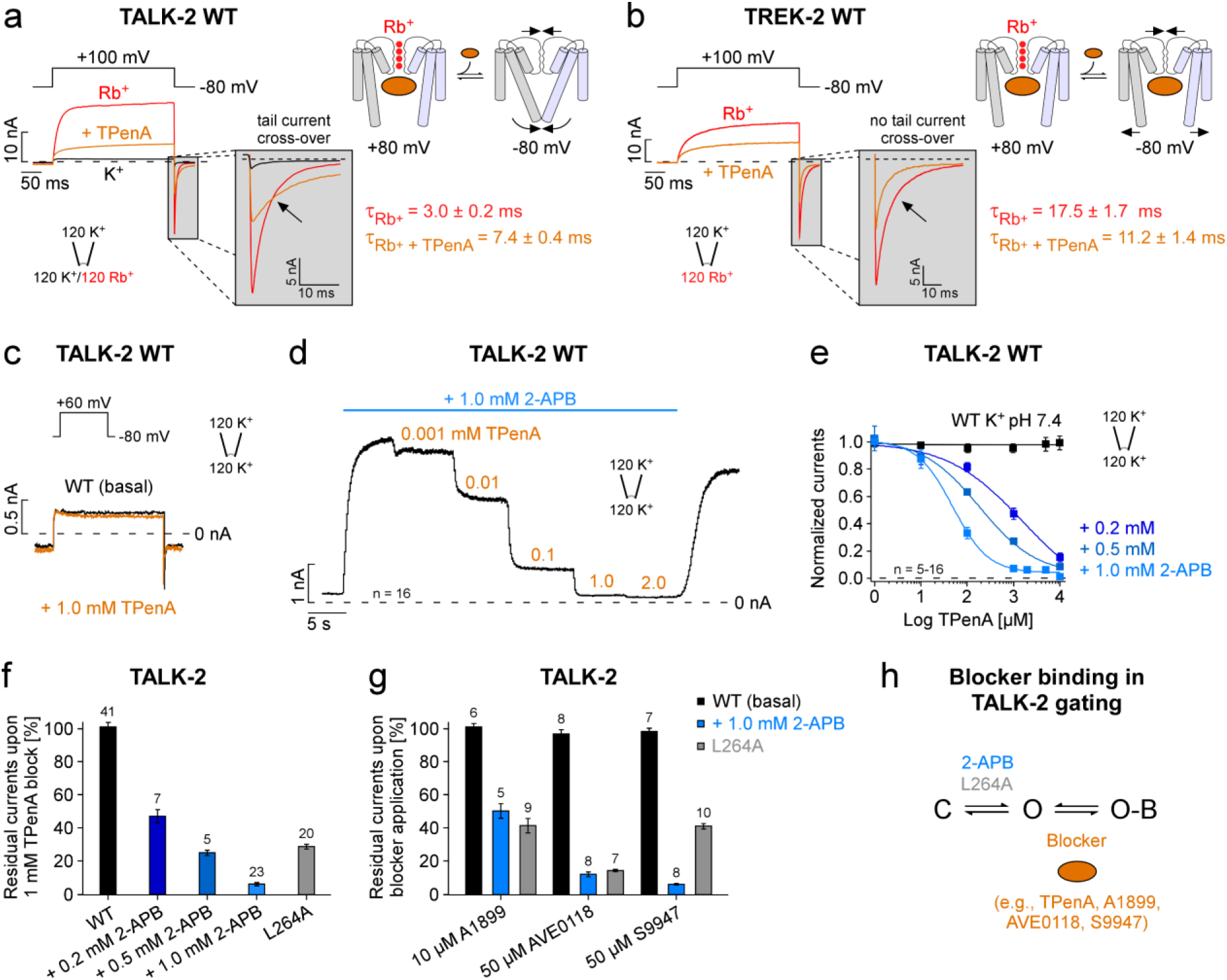
Open channel blocker show state-dependent pore accessibility and slowing of deactivation kinetics in TALK-2. **a** Current responses measured in inside-out patches under voltage-clamp conditions from *Xenopus laevis* oocytes expressing WT TALK-2 channels activated with indicated voltage steps under symmetrical ion conditions (120 mM [K^+^]_ex._/120 mM [X^+^]_int._) at pH 7.4 with either intracellular K^+^ (black trace, basal state) or Rb^+^ (red trace, activated state) or with 1.0 mM TPenA in Rb^+^ (orange trace). Note, the presence of TPenA shows slowing of deactivation resulting in a tail current cross-over (zoom-in). Cartoon depicting a simple model for TALK-2 channel gating, whereby Rb^+^ activation of the SF enables blocker (e.g., TPenA) binding in the pore cavity and unbinding facilitates lower and SF gate closure at −80 mV. **b** Same recording as in (a) with TREK-2 K_2P_ channels showing inhibition with 100 µM TPenA without tail current cross-over. **c** Representative current responses of WT TALK-2 channels to a 300 ms voltage step as indicated in the absence (black traces) and presence of 1.0 mM TPenA (orange traces) applied to the intracellular side of the membrane. **d** Representative measurement of TALK-2 channel currents at +40 mV showing dose-dependent TPenA inhibition in the pre-activated state with 1.0 mM 2-APB. **e** Dose-response curves of TPenA inhibition from measurements as in (d) for TALK-2 in unstimulated (basal) conditions (black) and pre-activated states with 2-APB (blue) with altering apparent affinities for TPenA (IC_50_ (0.2 mM 2-APB) = 778 ± 116, IC_50_ (0.5 mM 2-APB) = 215 ± 28, IC_50_ (1.0 mM 2-APB) = 54 ± 10). **f** Residual currents of WT and L264A mutant TALK-2 channels at +40 mV upon block of 1.0 mM TPenA at the indicated conditions. **g** Residual currents of unstimulated (black), 2-APB pre-activated WT (blue) and L264A mutant (gray) TALK-2 channels after inhibition with the indicated open channel blocker. **h** Simplified gating scheme indicating that blocker interact with the open state of TALK-2 to produce inhibition. Values are given as mean ± s.e.m with the number (n) of experiments indicated in the figure.

These findings suggest that TPenA is a state-dependent blocker in TALK-2 channels and, thereby, the apparent affinity for TPenA should strongly depend on the fraction of channels being open (**Fig. 5 h**). Indeed, in the basal state (low *Po*) hardly any TPenA block was observed (**Fig. 5c**), whereas 2-APB activation induced strong dose-dependent TPenA inhibition (**Fig. 5d-f**). Accordingly, the apparent affinity for TPenA inhibition increased dramatically with the degree of 2-APB activation (IC_50_ 0.2 mM 2-APB = 1108 ± 411 µM, IC_50_ 0.5 mM 2-APB = 225 ± 25 µM, IC_50_ 1.0 mM 2-APB = 54 ± 10 µM) (**Fig. 5e**). We further explored this observation by testing several other compounds known to block K_2P_ channels (e.g., TASK-1) such as A1899, AVE0118 and S9947^35^. Remarkably, for each blocker tested little inhibition was seen for unstimulated TALK-2 channels while strong inhibition was observed upon activation by 2-APB (**Fig. 5g**). Furthermore, the g-o-f mutation L264A resulted in TALK-2 channels with high sensitivity to inhibition for all tested blockers without 2-APB activation further indicating that the mutation promoted opening of the lower gate (**Fig. 5f, g, Supplementary Fig. 6a, b**).

In conclusion, the existence of the lower gate in TALK-2 transformed the state-independent pore block as seen in TREK-2 into a state-dependent (‘foot in the door’-like) block as typical for K_v_ channels.

### Opening of the lower gate reduces the mechanical load coupled to the voltage-powered SF gate

The voltage-dependent activation mechanism of the SF gate in K_2P_ channels allows to estimate the electrical work (ΔG = zFΔV_1/2_) required for pore opening by fitting the G-V curve to a Boltzmann equation. A leftward shift of the G-V curve (without a change in slope) indicates a reduction in the free energy between the closed and open state of the channel. The open state represents the situation of both gates being simultaneously open but voltage can only exert force on the SF gate that functions as voltage sensor. However, when the two gates are energetically coupled then any circumstance that opens the lower gate should also reduce the electrical work to open the SF gate (**Fig. 7d**). We tested this concept using TPenA that we have shown to hinder lower gate closure. Indeed, the presence of 1 mM TPenA caused a 46.1 ± 4.9 mV leftward shift of the G-V curve, while in TREK-2 K_2P_ channels exhibiting a constitutively open lower gate the G-V curve was not affected in the presence of TPenA (**Fig. 6a, b, Supplementary Table 4**). Likewise, 2-APB and LC-CoA that we have shown to open the lower gate also caused a leftward shift of the G-V curve (**Figure 6f, Supplementary Fig. 7a**). Further, stabilising the open state of SF directly by increasing pH_e_ also shifted the G-V curve leftwards, thus, deprotonation of the pH sensor reduces the free energy difference between the closed and open state of the SF (**Fig. 6d, e**). This implies that low pH and hyperpolarization induce a similar closed state of the SF. We further tested whether mutations that open the lower gate (L262A and L264A) would affect the extracellular pH sensitivity. Indeed, both mutations shifted the pH-current relationship towards more neutral pH suggesting that opening of the lower gate promoted the deprotonated pH sensor state indicating allosteric coupling of the lower gate to extracellular pH sensor (**Fig. 6g, Supplementary Fig. 7b**).

**Fig. 6.**
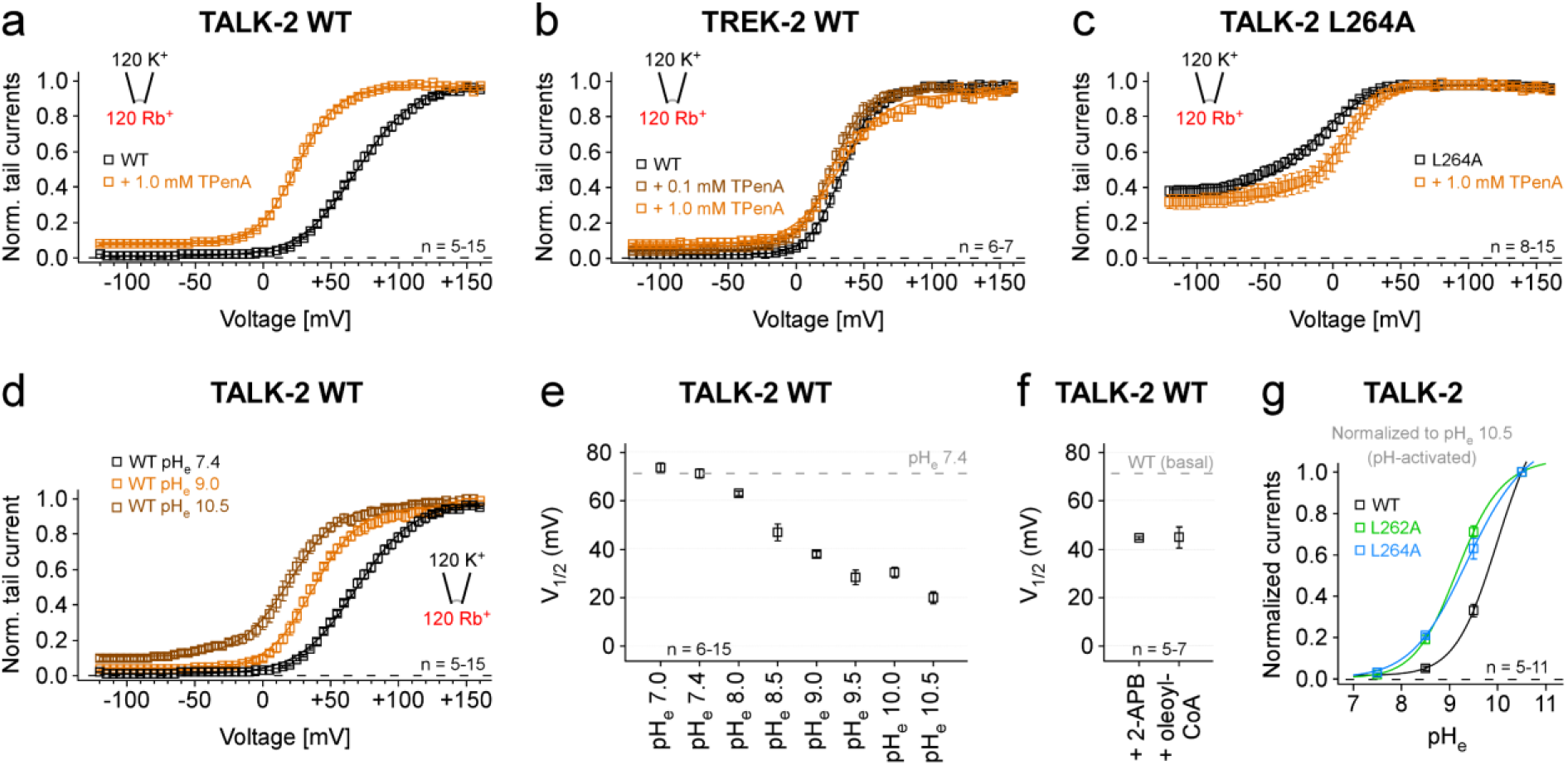
Impact of ligand modulation on SF energetics in TALK-2 K_2P_ channels. **a-c** G-V curves analyzed from current-voltage families (−120 mV to +160 mV with 5 mV increments) measured under symmetrical ion conditions with intracellular Rb^+^ of WT TALK-2 (a), WT TREK-2 (b) and L264A mutant TALK-2 channels (c) in the absence (black traces) and presence of 0.1 mM (brown trace) or 1.0 mM TPenA (orange traces), respectively. **d** G-V curves analyzed from WT TALK-2 tail currents in the presence of pH_e_ 7.4 (black trace, 9.0 (orange trace) and 10.5 (brown trace). **e** V_1/2_ values from G-V curves analyzed as in (d) with varying pH_e_ as indicated. **f** V_1/2_ values from G-V curves of WT TALK-2 channels activated with 1.0 mM 2-APB or 5.0 µM oleoyl-CoA. **g** Normalized currents from TEVC measurements of *Xenopus* oocytes expressing WT and mutant (L262A or L264A respectively) TALK-2 channels activated by increasing pH_e_ from 5.5 to 10.5 with 0.5 pH increments. Currents were elucidated with a voltage protocol ramped from −120 mV to +45 mV within 3.5 s, analyzed at +40 mV and normalized to pH 10.5. Values are given as mean ± s.e.m with the number (n) of experiments indicated in the figure and supplementary table 3 and 4 (n.d. = not determinable, n.-e. = non-expressing).

**Fig. 7.**
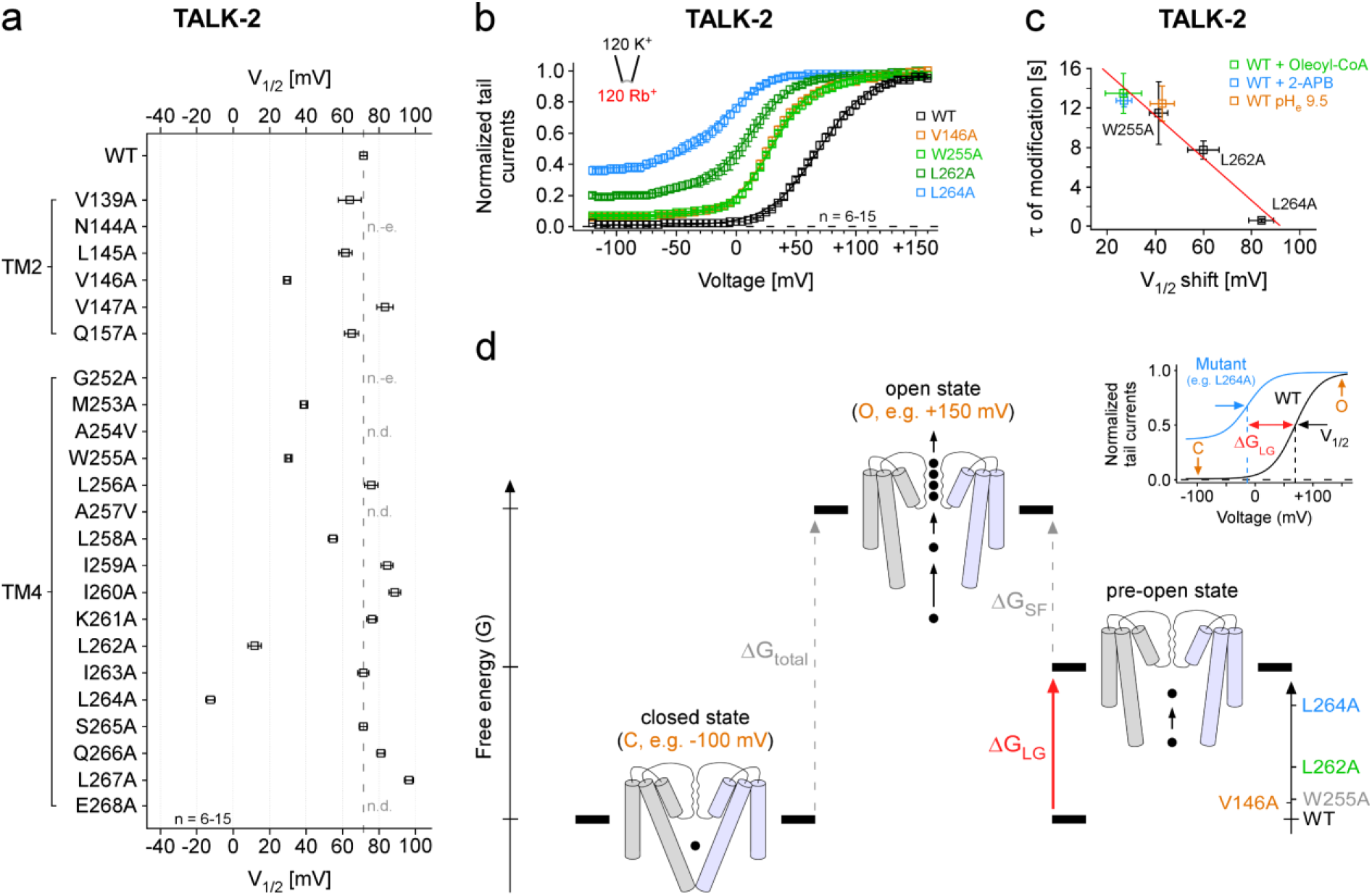
Functional coupling of the SF and the lower gate in TALK-2 K_2P_ channels. **a, b** G-V curves analyzed from tail currents at −80 mV after 300 ms pre-pulse steps (−120 mV to +160 mV with 5 mV increments) under symmetrical ion conditions with intracellular Rb^+^ of WT and mutant TALK-2 channels as indicated (b) and the summary of V_1/2_ values from Boltzmann fits to the corresponding G-V curves (a). **c** Correlation of the V_1/2_ shifts of mutant TALK-2 channels at basal and WT TALK-2 channels at indicated conditions with the time constants of modification of TALK-2 L145C channels under the corresponding activatory conditions or in combination with the respective g-o-f mutation. **d** Simplified energetic scheme depicting the electrical work (ΔG = zFΔV_1/2_) required to open both gates (ΔG_total_) with the individual contribution of the SF gate (ΔG_SF_) and lower gate (ΔG_LG_). Mutations (as indicated in the inlay) that open the lower gate reduced this electrical work as seen in the positive V_1/2_ shifts of the G-V curve. Note, our results actually show that both gates are strongly positively coupled and, thus, the pre-open state (just the lower gate open) is just a conceptual state to illustrate the energetic contribution of the lower gate. Values are given as mean ± s.e.m with the number (n) of experiments indicated in the figure and supplementary table 3 and 4 (n.d. = not determinable, n.-e. = non-expressing).

We further explored this concept of allosteric coupling using the mutations identified in the functional alanine screen of TM2 and TM4. Indeed, all mutations that produced a g-o-f effect also caused a leftward shift of the G-V curve (**Fig. 3a**, **Fig. 7a, b and Supplementary Table 3**). Further, the V_1/2_ shift was correlated to the increase in the L145C modification rate (**Fig. 7c**). Thus, the degree (i.e., frequency) of lower gate opening cause by the g-o-f mutations was correlated to the reduction in electric work to open the SF gate. Accordingly, the largest effect on the G-V curve (i.e., an 84.0 ± 5.3 mV shift) was seen for the L264A mutation that had the largest g-o-f effect, as well as caused the fastest modification of L145C and, thus, the highest *Po* of the lower gate (**Fig. 3a, k, Fig. 7a-c**). Actually, this mutation promoted the open state of the lower gate so strongly that 2-APB produced only a minor further current increase (**Supplementary Fig. 6c**), the speed of L145C MTS-ET modification at +40 mV was not further increased by Rb^+^ (**Supplementary Fig. 6d**) and TPenA has no marked effect on the G-V curve (**Fig. 6c**). Therefore, we conclude that the ∼84 mV shift in V_1/2_ produced by the L264A mutant might roughly represent the mechanical load of the conformational change that the voltage-powered SF gate has to move for opening the lower gate.

## Discussion

The present study on TALK-2 K_2P_ channels provides several lines of evidence suggesting the existence of a lower permeation gate at the cytoplasmic pore entrance in TALK-2. Firstly, access of cysteine modifying reagents to the inner pore cavity is blocked in closed TALK-2 channels, but possible in the presence of activating ligands, suggesting a pore entrance constriction. Furthermore, systematic alanine scanning mutagenesis identified several g-o-f mutations in distal parts of TM2 and TM4 that not only strongly activated TALK-2 but also removed the entrance constriction. Employing a TALK-2 homology models based on the TASK-1 and TASK-2 structures revealed that the g-o-f mutations cluster in a region corresponding to the lower gates identified in these channels. Finally, using Rb^+^ as a probe we show that the identified lower constriction is actually a permeation gate.

### Tight gate coupling defines the gating behavior in TALK-2 channels

In addition to the lower gate, TALK-2 channels are also operated by a SF gate similar to many other K_2P_ channels. We used three assays to dissect the status of the two gates and their coupling in the context of various gating stimuli: (i) the modification rate of L145C reports about the relative *Po* of the lower gate, (ii) the G-V curves report on the electrical work required to open the SF filter gate and (iii) the current amplitudes report on the fraction of channels with both gates open (overall channel *Po*). We found that the three parameters were strongly correlated in all g-o-f mutations suggesting a tight positive coupling of the two gates (**Fig. 7d**). This concept is also supported by a number of additional findings reported here. Opening the SF by high pH_e_ or voltage, likewise, opened the lower gate (i.e., increased L145C modification rate). Reciprocally, ligands that opened the lower gate (i.e., increased the rate of L145C modification) such as 2-APB or LC-CoA also positively shifted the G-V curve (opening of the SF gate). Particular striking was the observation that exchange of the permeating ion from K^+^ to Rb^+^ caused a large increase in the L145C modification rate indicating that the mere difference in SF ion occupancy is sufficient to induce a marked structural change at the pore entrance (i.e., opening the lower gate). In structural terms the strong positive gate coupling can be envisioned as rigid connection of the two gates within the protein structure and, thus, opening/closing of one gate also forces the other gate to open/close.

### How is the lower gate coupled to the selectivity filter?

We speculate that movement of TM4 underlies lower gate opening as well as force transduction onto the SF gate to induce its opening in TALK-2. Thus, the TM4 might represent the rigid connection linking the two gates responsible for the tight gate coupling. This hypothesis appears appealing as a similar mechanism has been shown to couple the lower gate to the SF gate in MthK (prokaryotic K^+^ channel from *Methanobacterium thermoautotrophicum*), based on MD simulations performed with different opening diameters at the lower gate^36^. Here an isoleucine (I84) at the end of TM2 (corresponding to TM4 in TALK-2) was shown to exert force on a threonine (T59) of the SF to promote the conductive SF state. Although members of the TREK/TRAAK subfamily lack a functional lower gate^28,29^, the TM4 also moves during temperature-, mechano- or lipid-induced gating (known as the down- to up-state transition^37,38^) and this movement is also thought to open the SF gate^25,26^. Furthermore, also the lower ‘X-gate’ in TASK-1 is formed by TM4 and, thus, envisioned to move during gating^31^. Therefore, movement of the TM4 segments coupled to the SF gate might be the general theme in K_2P_ channel gating (as well as other K^+^ channels). In some channels this movement may result in the formation of a lower gate (e.g., in TASK-1, TASK-2 and TALK-2) and in other channels not (e.g., in TREK-1/-2 and TRAAK), while ultimately the SF is affected in the activation process. How this coupling is realized in atomic detail in TALK-2 warrant future studies and in particular high resolution structural information of different states.

### Implications of the state-dependent pore access for TALK-2 gating and pharmacology

Functionally, the lower gate in TALK-2 resembles the helix-bundle crossing gate in voltage-gated K_v_ channels. In K_v_ channels voltage is thought to open the helix-bundle crossing gate via the voltage-dependent movement of segment 4 (S4)^39,40^. By marked contrast, in K_2P_ channels, voltage opens the SF gate via a voltage-dependent ion binding step^20^ that forces the filter in its conductive state. In TALK-2 K_2P_ channels this structural change in the SF appears also to force the lower gate open as apparent in voltage-dependent modification rate of L145C. Upon repolarization, the ion-flux inversion is thought to change the SF ion occupancy leading to its inactivation as visible in the fast reduction of the tail current amplitudes^20^. Intriguingly, in the presence of the pore blocker TPenA the decay of the current is slowed resembling the ‘foot in the door’ effect seen in K_v_ channels^33,34^. This suggest that blocker binding to the pore cavity prevented the lower gate from closing. Importantly, this also suggest that TPenA concurrently prevented the SF gate from closing as blocker unbinding from a channel with a non-conductive SF would be electrophysiological invisible and, thus, would not produce the apparent slowing of deactivation. In agreement, in TREK-2 channels (lacking the lower gate) TPenA had no effect on the tail current kinetics indicating that the SF gate can close unhindered with the blocker bound upon inversion of the electric field upon repolarization. What stops now the SF gate from closing with TPenA bound in TALK-2? In K_v_ channels blocker binding to the pore cavity prevents lower gate closure (the actual structural change is still unknown) suggesting that the pore cavity changes its structure (and possible its size) in concert with the lower gate closure and this change is hindered (or delayed) by the blocker^41,42^. Thus, we presume that a similar structural change is also occurring in TALK-2 and is also prevented by TPenA. Consequently, because the SF gate and the lower gate are tightly coupled, TPenA also disturbs the SF gate in the closing reaction. In further agreement, we found that TPenA caused a leftward shift of the G-V curve as expected for a blocker that only binds to the pore with both gates open.

In single channel recordings we estimated the basal activity (i.e., the *NPo*) for WT TALK-2 to about 4 % and, thus, only a small fraction of channels would be sensitive to a pore blocker if only open channels can bind the compound (**Fig. 5h**). Indeed, we observed very little current inhibition when TPenA was applied to TALK-2 channels in absence of an activating stimulus, but strong inhibition was observed for activated TALK-2 channels. This behavior was also seen for various other small molecule pore blocker, thus, state-dependent pore inhibition appears to be a defining feature of the TALK-2 channel pharmacology as this was so far not seen in any other K_2P_ channel.

### Gate coupling in other K^+^ channels: Differences and similarities

The established gating cycles of K_v_ and KcsA (prokaryotic K^+^ channel from the soil bacterium *Streptomyces lividans*) channels suggest that the activation (lower) gate serves as the primary ion permeation barrier that needs to open to allow current flow, while in a following step the SF inactivates to terminate ion permeation^39,40,43,44^. Thus, in these cases the two gates are coupled sequentially and, in a negative manner. However, it is currently unknown whether the SF is open (conductive) or closed (non-conductive) in the resting (not activated) state of K_v_ or KcsA channels because it is not possible to directly measure the conductivity of the SF in a closed channel. In TALK-2 K_2P_ channels the SF serves as a gate as well as the voltage sensor. The latter property provides information on stability of the conductive state of the filter that can be extracted from the V_1/2_ value of the corresponding G-V curve. Here, we show that when the lower gate is mostly closed then it requires a large amount of electrical work to open the SF, i.e., a membrane depolarization to 72 ± 2 mV (V_1/2_) for half maximal voltage activation. However, when the lower gate is mostly open as seen in the L264A TALK-2 mutant channel this work is strongly reduced (V_1/2_ = −12 ± 2 mV) and, thus, might approximately reflect the work required to open the lower gate. Actually, even very negative potentials cannot close the SF gate to a large degree as seen by the pedestal for the relative *Po* that levels off at ∼0.4 in TALK-2 L264A channels (**Fig. 7b**). Notably, these results closely resemble the constitutively open phenotype in *Shaker* K^+^ channels seen with mutations at the assumed hydrophobic seal of the helix-bundle crossing gate. These mutations also result in strong positively shifted G-V curves and high pedestal *P_O_*s even at very negative potentials^45^. Interestingly, the two residues with the strongest g-o-f effect (V146A and L264A) in TALK-2 are also hydrophobic and in direct proximity (according to our TASK-1 based homology model) and, thus, might also form a hydrophobic seal^46^ that, thereby, could represent the lower gate in TALK-2.

Intriguingly, a recent MD simulation study on KcsA suggest that the SF might be non-conductive when the lower (activation) gate is closed as this also implies positive coupling of the two gates in this archetypical K^+^ channel^47^. Here, using electrophysiological means, we have demonstrated directly that the SF gate in TALK-2 K_2P_ channels opens and closes in concerts with the lower gate. It will be interesting to see whether this concept also applies to other K^+^ channels possessing two gates such as members of the large family of voltage-gated K^+^ channels.

## Methods

### Molecular biology

In this study the coding sequences of human K_2P_2.1 TREK-1 (GenBank accession number: NM_172042), human K_2P_10.1 TREK-2 (NM_021161) and human K_2P_17.1 TALK-2 (EU978944.1)/human K_2P_17.1 TASK-4 (NM_031460.3) were used.

For K^+^ channel constructs expressed in *Xenopus laevis* oocytes the respective K^+^ channel subtype coding sequences were subcloned into the oocyte expression vector pSGEM or the dual-purpose vector pFAW which can be used for HEK293 cell expression as well and verified by sequencing. All mutant channels were obtained by site-directed mutagenesis with custom oligonucleotides. Vector DNA was linearized with NheI or MluI and cRNA synthesized *in vitro* using the SP6 or T7 AmpliCap Max High Yield Message Maker Kit (Cellscript, USA) or HiScribe® T7 ARCA mRNA Kit (New England Biolabs) and stored at −20°C (for frequent use) and −80 °C (for long term storage).

### Electrophysiological recordings in *Xenopus laevis* oocytes

#### Two-electrode voltage-clamp (TEVC) measurements

Electrophysiological studies were performed using the TEVC technique in *Xenopus laevis* oocytes. Ovarian lobes were obtained from frogs anesthetized with tricaine. Lobes were treated with collagenase (2 mg/ml, Worthington, type II) in OR2 solution containing (in mM): 82.5 NaCl, 2 KCl, 1 MgCl_2_, 5 HEPES (pH 7.4 adjusted with (NaOH/HCl) for 2 h. Isolated oocytes were stored at 18 °C in ND96 recording solution (in mM): 96 NaCl, 2 KCl, 1.8 CaCl_2_, 1 MgCl_2_, 5 HEPES (pH 7.5 adjusted with NaOH/HCl) supplemented with Na-pyruvate (275 mg/l), theophylline (90 mg/l), and gentamicin (50 mg/l). Oocytes were injected with 50 nl of cRNA for WT or mutant TALK-2 and incubated for 2 days at 18 °C. Standard TEVC measurements were performed at room temperature (21 - 22 °C) with an Axoclamp 900A amplifier, Digidata 1440A, and pClamp10 software (Axon Instruments, Molecular Devices, LLC, USA). Microelectrodes were fabricated from glass pipettes, back-filled with 3 M KCl, and had a resistance of 0.2 - 1.0 MΩ.

#### Inside-out patch-clamp measurements

*Xenopus laevis* oocytes were surgically removed from anesthetized adult females, treated with type II collagenase (Sigma-Aldrich/Merck, Germany) and manually defolliculated. 50 nl of a solution containing the K^+^ channel specific cRNA was injected into Dumont stage V - VI oocytes and subsequently incubated at 17 °C in a solution containing (mM): 54 NaCl, 30 KCl, 2.4 NaHCO_3_, 0.82 MgSO_4_ x 7 H_2_O, 0.41 CaCl_2_, 0.33 Ca(NO_3_)_2_ x 4 H_2_O and 7.5 TRIS (pH 7.4 adjusted with NaOH/HCl) for 1 - 7 days before use. Electrophysiological recordings: Excised patch measurements in inside-out configuration under voltage-clamp conditions were performed at room temperature (22 - 24 °C). Patch pipettes were made from thick-walled borosilicate glass GB 200TF-8P (Science Products, Germany), had resistances of 0.2 - 0.5 MΩ (tip diameter of 10 - 25 µm) and filled with a pipette solution (in mM): 120 KCl, 10 HEPES and 3.6 CaCl_2_ (pH 7.4 adjusted with KOH/HCl). Intracellular bath solutions and compounds were applied to the cytoplasmic side of excised macro patches for the various K^+^ channels via a gravity flow multi-barrel pipette system. Intracellular solution had the following composition (in mM): 120 KCl, 10 HEPES, 2 EGTA and 1 Pyrophosphate (pH adjusted with KOH/HCl). These patch pipette and intracellular solutions were used for recordings of K_2P_, K_v_ and hERG channels (pH 8.0 for measurements of TREK-1/-2 WT and mutant channels to avoid pH activation; otherwise, pH 7.4). In other intracellular bath solutions, K^+^ was replaced by Rb^+^ (pH 7.4 adjusted with RbOH/HCl). Currents were recorded with an EPC10 amplifier (HEKA electronics, Germany) and sampled at 10 kHz or higher and filtered with 3 kHz (−3 dB) or higher as appropriate for sampling rate.

#### Inside-out single channel patch-clamp measurements

Single channel patch clamp measurements in the inside-out configuration under voltage-clamp conditions using *Xenopus laevis* oocytes were performed similar as previously described ^48^. Briefly, the vitelline membranes of the oocytes were manually removed after shrinkage by adding mannitol to the bath solution. All experiments were conducted at room temperature (21 - 22 °C) 1 - 2 days after injection of 50 nl TALK-2 cRNA. Borosilicate glass capillaries GB 150TF-8P (Science Products, Germany) were pulled with a DMZ-Universal Puller (Zeitz Instruments, Germany) and had a resistance of 4 - 6 MΩ when filled with pipette solution containing (in mM): 120 KCl, 10 HEPES, 3.6 CaCl_2_ (pH 8.5 adjusted with KOH/HCl). Bath solution had the following composition (in mM): 120 KCl, 10 HEPES, 2 EGTA, 1 Pyrophosphate (pH 8.5 adjusted with KOH/HCl). Gap-free voltage pulses of −100 mV were continuously applied. Single channel currents were recorded with an Axopatch 200B amplifier, a Digidata 1550B A/D converter and pClamp10 software (Axon Instruments, Molecular Devices, LLC, USA) and were sampled at 15 kHz with the analogue filter set to 5 kHz. Additionally, data was digitally filtered by 2 kHz (Lowpass, Bessel) with ClampFit10 before analysis. Data were analyzed with ClampFit10 and Origin 2016 (OriginLab Corporation, USA). The single channel search tool of the ClampFit10 software was used to identify and analyze the single channel events in a time frame of 60 s. A group of single channel events (minimum 3 events) has been classified as a burst, if the time between two single channel openings were lower than five times of the short-closed time (5 x t_C1_).

### Animals

The investigation conforms to the guide for the Care and Use of laboratory Animals (NIH Publication 85-23). For this study, twenty-five female *Xenopus laevis* animals were used to isolate oocytes. Experiments using Xenopus toads were approved by the local ethics commission.

### Drugs, chemical compounds and bioactive lipids

Tetra-pentyl-ammonium chloride (TPenA), 2-aminoethoxydiphenyl borate (2-APB) (Sigma-Aldrich/Merck, Germany), BL-1249, A1899 (Tocris Bioscience, Germany), AVE0118, S9947 (Axon Medchem, Germany) and oleoyl-CoA (LC-CoA 18:1) (Avanti Polar Lipids, USA) were prepared as stocks (1 - 100 mM) in DMSO, stored at −80 °C and diluted to the final concentration in the intracellular recording solution. (2-(Trimethylammonium)ethyl) MethaneThioSulfonate Chloride (MTS-ET) (Toronto Research Chemicals, USA) was directly dissolved to the desired concentration of 1.0 mM in the intracellular recording solution prior to each experiment. MTS-ET was used immediately after dilution for maximally 5 min.

### Homology modelling - TALK-2 models based on TASK-1 or TASK-2

Human TASK-1 crystallographic structure (PDB ID: 6RV3, chains A, B)^31^ and mouse TASK-2 cryo-electron microscopy (Cryo-EM) structure (PDB ID: 6WLV, chains A, B)^30^ were used as templates to build the TALK-2 models. First, the structures were prepared using ‘Protein preparation wizard’ module of the Schrödinger’s suite software (Protein Preparation Wizard; Epik, Schrödinger, LLC, New York, NY, 2018-4; Impact, Schrödinger, LLC, New York, NY, 2018-4; Prime, Schrödinger, LLC, New York, NY, 2018-4). Charges and parameters were assigned according to the force field OPLS-2005^49^. The missing residues in TASK-1 (149 - 151 from chain A and 150 - 151 from chain B) were modeled using ‘crosslink protein’ tool from the Schrödinger suite.

The sequence of TALK-2 (KCNK17) protein was obtained from the GenBank database^50^. Alignments between the template protein (TASK-1 or TASK-2) sequences and the TALK-2 sequence were performed using the Smith-Waterman algorithm^51^ with BLOSUM62^52^ matrix for scoring the alignment. The modeling was carried out using the BioLuminate^53^ module of the Schrödinger suite. The standard modeling protocol was used, which consists of a search for rotamers for non-conserved residues and loops to eliminate clashes and then an energetic minimization with the OPLS2005 force field in vacuum. The protonation states of the TALK-2 models were predicted at a pH of 7.0 with PROPKA^54^, and another energetic minimization of the hydrogen atoms was performed using the conjugate gradient method^55^ implemented in the Schrödinger suite. The sequence of the TALK-2 model based on 6RV3 extends from residue R15 to K282 and based on 6WLV from residue T22 to H278, respectively. It was verified with the procheck^56^ software that 91.2 % and 93,5 % of the residues are in the most favored regions for the models based on 6RV3 and 6WLV, respectively.

Alignments:

6RV3:

MKRQNVRTLALIVCTFTYLLVGAAVFDALESEPELIERQRLELRQQELRARYN∼LSQGGYEELERVVLRLKPHKAGVQ∼∼∼∼∼∼∼∼WRFAGSFYFAITVITTIGYGHAAPSTDGGKVFCMFYALLGIPLTLVMFQSLGERINTLVRYLLHRAKKGLGMRRADVSMANMVLIGFFSCISTLCIGAAAFSHYEHWTFFQAYYYCFITLTTIGFGDYVALQKDQALQTQPQYVAFSFVYILTGLTVIGAFLNLVVLRFMTMNAEDEKRDAENL

TASK4 (TALK-2):

RGCAVPSTVLLLLAYLAYLALGTGVFWTLEGRAAQDSSRSFQRDKWELLQNFTCLDRPALDSLIRDVVQAYKNGASLLSNTTSMGRWELVGSFFFSVSTITTIGYGNLSPNTMAARLFCIFFALVGIPLNLVVLNRLGHLMQQGVNHWASRLGGTWQDPDKARWLAGSGAL∼LSGLLLFLLLPPLLFSHMEGWSYTEGFYFAFITLSTVGFGDYVIG∼MNPSQRYPLWYKNMVSLWILFGMAWLALIIKLILSQLETPGRVCSCCHHSSK

6WLV:

GPLLTSAIIFYLAIGAAIFEVLEEPHWKEAKKNYYTQKLHLLKEFPCLSQEGLDKILQVVSDAADQGVAITGNQT∼FNNWNWPNAMIFAATVITTIGYGNVAPKTPAGRLFCVFYGLFGVPLCLTWISALGKFFGGRAKRLGQFLTRRGVSLRKAQITCTAIFIVWGVLVHLVIPPFVFMVTEEWNYIEGLYYSFITISTIGFGDFVAGVNPSANYHALYRYFVELWIYLGLAWLSLFVNWKVSMFVEVHKAIKKRR

TASK4 (TALK-2):

TVLLLLAYLAYLALGTGVFWTLEGRAAQDSSRSFQRDKWELLQNFTCLDRPALDSLIRDVVQAYKNGASLLSNTTSMGRWELVGSFFFSVSTITTIGYGNLSPNTMAARLFCIFFALVGIPLNLVVLNRLGHLMQQGVNHWASRLGGTWQDPDKARWLAGSGALLSGLLLFLLLPPLLFSHMEGWSYTEGFYFAFITLSTVGFGDYVIGMNPSQRYPLWYKNMVSLWILFGMAWLALIIKLILSQLETPGRVCSCCH

### Data acquisition and statistical analysis

Data analysis and statistics were done using Fitmaster (HEKA electronics, version: v2×73.5, Germany), Microsoft Excel 2021 (Microsoft Corporation, USA) and Igor Pro 9 software (WaveMetrics Inc., USA).

Recorded currents were analyzed from stable membrane patches at a voltage defined in the respective figure legend or with a voltage protocol as indicated in the respective figure. The fold activation (fold change in (tail) current amplitude) of a ligand (drug or bioactive lipid) was calculated from the following equation:

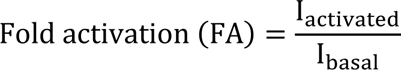

with I_activated_ represents the stable current level in the presence of a given concentration of a respective ligand and I_basal_ the measured current before ligand application. Percentage inhibition or residual currents upon blocker application for a ligand (drug or bioactive lipid) was calculated from stable currents of excised membrane patches using the following equation below:

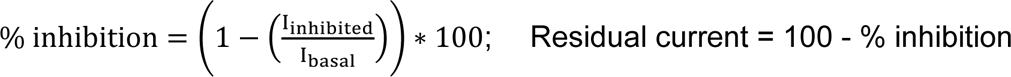

where I_inhibited_ refers to the stable current level recorded in the presence of a given concentration of a drug or bioactive lipid and I_basal_ to the measured current before ligand application. The macroscopic half-maximal concentration-inhibition relationship of a ligand was obtained using a Hill-fit for dose-response curves as depicted below:

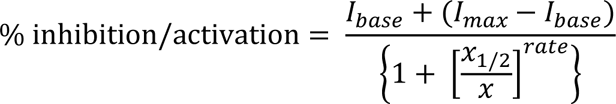

with base and max are the currents in the absence and presence of a respective ligand, x is the concentration of the ligand, x_1/2_ is the ligand concentration at which the activatory or inhibitory effect is half-maximal, rate is the Hill coefficient.

For analysis of activation and deactivation time constants (τ) as well as the time constants of MTS modification (τ) current traces were fitted with a mono-exponential equation as depicted below:

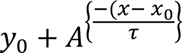

Conductance-voltage (G-V) relationships were determined from tail currents recorded at a holding potential (V_H_) of −80 mV after 300 ms depolarizing steps as indicated in the respective figure legend. Data were analyzed with a single Boltzmann fit following the equation below:

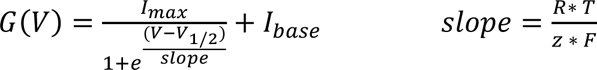

with V_1/2_ represents the voltage of half-maximal activation, s is the slope factor and I_max_ and I_base_ represent the upper and lower asymptotes. Under appropriate experimental conditions, you can use slope to calculate the valence (charge) of the ion moving across the channel. Slope equals R * T/ z * F where R is the universal gas constant, T is temperature in K, F is the Faraday constant, and z is the valence.

Data from individual measurements were normalized and fitted independently to facilitate averaging.

Throughout the manuscript all values are represented as mean ± s.e.m. with n indicating the number of individual executed experiments. Error bars in all figures represent s.e.m. values with numbers (n) above indicating the definite number of executed experiments. A Shapiro-Wilk test or Kolmogorow-Smirnow test was used to determine whether measurements were normally distributed. Statistical significance between two groups (respective datasets) was validated using an unpaired Student’s *t*-test or Wilcoxon rank test after *f*-test application. Asterisks indicate the following significance: * *P* ≤ 0.05, ** *P* ≤ 0.01 and *** *P* ≤ 0.001. Zero current level was indicated using dotted lines in all figures.

Image processing and figure design was done using Igor Pro 9 (64 bit) (WaveMetrics, Inc., USA) and Canvas X Draw (Version 20 Build 544) (ACD Systems, Canada).

## Data availability

Data supporting the findings of this manuscript are available from the corresponding authors upon request. This study includes no data deposited in external repositories.

## Acknowledgments

We thank the members of our laboratories for technical support and helpful comments on the manuscript. These studies were supported by the Deutsche Forschungsgemeinschaft (DFG, German Research Foundation) to N.D. (DE1482-9/1) and T.B. (BA 1793/6-2)/ M.S. (SCHE 2112/1-2) as part of the Research Unit FOR2518, *DynIon*.

## Author contributions

N.D., T.B. and M.S. conceived and supervised the project; L.C.N., E.B.R., B.C.J., J.L., B.E. and M.S. performed patch-clamp experiments and analyzed the data. S.R. and N.D. planed and generated mutant constructs for two-electrode voltage-clamp experiments. F.-R.S. and S.R. performed and A.K.K. and S.R analyzed TEVC experiments. N.D. planned, S.R. performed and A.K.K. analyzed single channel recordings. S.C. planned mutant constructs for patch-clamp experiments. M. B. generated the TALK-2 homology model. M.S. prepared figures; T.B. and M.S. wrote the article with contributions from all authors.

